# Kinetic characterization of three human DExD/H-box RNA helicases

**DOI:** 10.1101/2025.02.07.637080

**Authors:** Fengling Li, U Hang Chan, Julia Garcia Perez, Hong Zeng, Irene Chau, Yanjun Li, Alma Seitova, Levon Halabelian

## Abstract

Human DExD/H-box RNA helicases are ubiquitous molecular motors that unwind and rearrange RNA secondary structures in an ATP-dependent manner. These enzymes play essential roles in nearly all aspects of RNA metabolism. While their biological functions are well-characterized, the kinetic mechanisms remain relatively understudied *in vitro*. In this study, we describe the development and optimization of a bioluminescence-based assay to kinetically characterize three human RNA helicases: MDA5, LGP2, and DDX1. The assays were conducted using annealed 24-mer RNA (blunt-ended double-stranded RNA) or double-stranded RNA (ds-RNA) with a 25- nt 3ʹ overhang. These findings establish a robust and high-throughput *in vitro* assay suitable for a 384-well format, enabling the discovery and characterization of inhibitors targeting MDA5, LGP2, and DDX1. This work provides a valuable resource for advancing our understanding of these helicases and their therapeutic potential in Alzheimer’s disease.

## Introduction

Helicases are enzymes that unwind nucleic acids, which is a fundamental requirement for many cellular metabolic processes across viruses, bacteria, and eukaryotes ^1^. Mutations in human helicase genes have been implicated in numerous human diseases, including cancer and neurodegeneration ^2^. Helicases are classified into six superfamilies (SF1-6) based on conserved sequence motifs and structural features. Monomeric helicases belong to superfamilies 1 and 2, while oligomeric ring-forming helicases are categorized in superfamilies 3 to 6 ^3^. Other studies, however, classify RNA helicases into five superfamilies (SF1-5) ^4–5^. Despite these variations in classification, most RNA helicases are part of the SF2 superfamily, which includes DExD/H-box helicases. These helicases are characterized by conserved DEAD (Asp-Glu-Ala-Asp), DExH (Asp-Glu-x-His), DExD (Asp-Glu-x-Asp), and DEAH (Asp-Glu-Ala-His) motifs in their ATP-binding domains ^5^.

DExD/H-box RNA helicases typically function as part of multi-protein complexes, with additional ATP-independent roles arising from interactions with protein partners ^6^. This versatility enables DExD/H RNA helicases to regulate various aspects of RNA metabolism ^3^. One important subgroup of helicases is the RIG-I-like receptor (RLRs) family, which plays a crucial role in the immune response to viral RNA.^7^ The RLR family consists of three members: retinoic acid-inducible gene I (RIG-I), melanoma differentiation-associated gene 5 (MDA5 or IFIH1), and laboratory of genetics and physiology 2 (LGP2 or DHX58) ^8^. RIG-I was the first identified, and is the most extensively studied member of this family, though it will not be discussed further here ^9^.

MDA5 is a cytoplasmic RNA helicase that functions as a pattern recognition receptor (PRR) in the innate immune system, particularly in the detection of viral RNA^10^. Upon binding to viral RNA, MDA5 activates downstream signaling pathways, leading to the production of type I interferons and other pro-inflammatory cytokines. Working alongside RIG-I, MDA5 plays a crucial role in antiviral immune responses ^11^. Mutations in MDA5 have been linked to autoimmune diseases such as Aicardi-Goutières syndrome and systemic lupus erythematosus^12^, underscoring its importance in immune regulation and autoimmune pathogenesis.

LGP2, another member of the RLR family, regulates both MDA5- and RIG-I-mediated antiviral responses ^13^. Unlike MDA5 and RIG-I, LGP2 lacks the caspase activation and recruitment domains required for signaling ^14^. Instead, it modulates the activity of these receptors and plays a regulatory role in antiviral immune responses. LGP2 is also implicated in other cellular processes, such as RNA stability and translation regulation, although its precise functions in these contexts remain under investigation ^14–15^ .

In contrast, non-RLR helicases from the DEAD-box family of DExD/H-box RNA helicases are distinguished by two RecA-like domains, RecA1 and RecA2. These DEAD-box helicases participate in various cellular processes, such as RNA processing, translation, and repair, rather than direct immune responses ^16^. Non-RLR helicases, such as DDX1, fall into this category ^17^. DDX1 is involved in various aspects of RNA metabolism, including RNA splicing, translation, and mRNA export ^18^. It also plays a role in viral infection by interacting with viral RNA and modulating host antiviral responses ^19^. DDX1 has also been implicated in cancer, contributing to tumor progression and metastasis through its effects on RNA metabolism and gene expression ^17^. Additionally, DDX1 interacts with NF-kappaB, a key regulator of immune response ^20^. Moore *et al* ^21^ demonstrated that DDX1 catalyzes the hydrolysis of ATP and deoxy-ATP in the presence of RNA, with its activity enhanced by various substrates, including single-stranded RNA molecules, a blunt-ended double-stranded RNA molecule, a hybrid of a double-stranded DNA-RNA molecule, and a single-stranded DNA molecule, as assessed by Thin Layer Chromatography (TLC) assays ^21^.

Overall, human DExD/H-box RNA helicases play critical roles in immune responses, viral defense mechanisms, and cellular homeostasis, making them important targets for research in infectious diseases, autoimmune disorders, and cancer biology. Additionally, the TREAT-AD (TaRget Enablement to Accelerate Therapy development for Alzheimer’s Disease) Center has identified MDA5, LGP2, and DDX1 as potential risk factors in Alzheimer’s disease progression 22.

Despite their biological significance, comprehensive kinetic characterization of human RNA helicases remains limited. In this study, we report the kinetic characterization of three DExD/H-box RNA helicases conducted in parallel in the presence of 24-mer RNA or ds-RNA.

## Materials and Methods

### Reagents

Kinase-Glo reagents (Cat# V6712: Luminescent signal linear to 10μM and Cat# V3772: Luminescent signal linear to 100μM) were purchased from Promega (Madison, WI, USA). ATP disodium salt (Cat# A2383) was obtained from Sigma-Aldrich (St. Louis, MO, USA). The white 384-well Plates (Item No.: 781207) were obtained from Greiner Bio-One North America Inc. (North Carolina, USA). ssDNA (30-mer of poly T), 24-mer of RNA (GGGACGUCAUGCGCAUGACGUCCC), and 22-bp double-stranded RNA (ds-RNA) with a 25-nt 3ʹ overhang (Strand 1: 5ʹ- UCGUGGCAUUUCUGCGUCGUUCUUUUCUUUUCUUUUCUUUUCUUUUC-3ʹ, Strand 2: 5ʹ-GAACGACGCAGAAAUGCCACGA-3ʹ) were ordered from IDT (Integrated DNA Technologies, USA). The 24-mer RNA was treated with an annealing program (heated to a high temperature 95°C for 2 minutes, and then slowly cooled down to room temperature) prior to use.

### Expressions and Purifications of DExD/H-box RNA helicases

The wild-type genes DDX1 (1-740 aa), MDA5A (306-1025 aa), MDA5A-FL (1-1025 aa), MDA5A-S (306-890 aa), and LGP2 (1-678 aa) were PCR-amplified and subcloned into the appropriate vectors: pFBOH-MHL for DDX1, pFBD-BirA for MDA5A, MDA5A-FL, and LGP2, and pET28-MHL for MDA5A-S. The resulting plasmids for DDX1, MDA5A, MDA5A- FL, and LGP2 transformed into DH10Bac™ Competent E. coli (Invitrogen) and a recombinant viral bacmid DNA was purified and followed by a recombinant baculovirus generation for baculovirus mediated protein production in *Sf9* insect cells^23^. The His6-tagged MDA5A-S (306- 890 aa) was expressed in *E. coli* BL21 (DE3) codon plus RIL strain (Stratagene), induced overnight at 15°C with 1 mM isopropyl-β-D-thiogalactopyranoside (IPTG).

After harvesting, cells were resuspended in lysis buffers: 20 mM Tris-HCl, pH 8, 500 mM NaCl, 5 mM imidazole, 5% glycerol, and 1X protease inhibitor cocktail for DDX1 and LGP2, and 20 mM Hepes, pH 7.5, 500 mM NaCl, 5 mM imidazole, 5% glycerol, and 1X protease inhibitor cocktail for MDA5A. Cells were lysed by rotating with NP40 (final concentration 0.6%) for 30 minutes, in the presence of benzonase nuclease, followed by sonication at a frequency of 7 (5” on/7” off) for 2 minutes (Sonicator 3000, Misonix). The crude lysates were clarified by high-speed centrifugation (60 minutes at 14,000 rpm at 10°C). The clarified lysate was loaded onto Hispur™ Ni-NTA resin (Thermo Scientific, Waltham, MA) and further purified by gel filtration on a Superdex 200 column (26 × 60) using an ÄKTA FPLC system (GE Healthcare, Little Chalfont, UK). Protein purity and quality were assessed and confirmed using SDS-PAGE and LC-MS. Pure fractions were pooled, concentrated, and flash-frozen.

### ATPase Assay for Determining Nucleotide Specificity

The ATPase activities of three helicases were monitored in *vitro* by a bioluminescent assay. Measurement of ATP depletion employed the Kinase-Glo assay system where a firefly luciferase detection reagent containing d-luciferin and buffer components are added to detect the remaining ATP following the helicase assays ^24–25^. For the experiments, ATP alone or ATP combined with different nucleotides (24-mer RNA, ds-RNA, or ssDNA) were used as substrates for the ATPase activities of the three helicases, respectively. The reaction mixtures contained 2 µM ATP, 100 nM of ssDNA or 24-mer RNA, or 3 nM ds-RNA in 20 mM Tris-HCl buffer (pH 7.5, 0.01% Tween 20, 1 mM MgCl₂, and 2 mM DTT), with a final volume of 15 µL per well. The reactions were initiated by adding helicases at final concentrations of 20 nM for MDA5, and 100 nM for DDX1 or LGP2. The reaction mixtures were incubated for 30-60 minutes, and then quenched by adding 15 µL of luciferase reagent mix each well. Subsequently, 25 µL of the mixture from each well was transferred to a 384-well white detection plate and incubated for an additional 15 minutes before measuring the generated signals using a BioTek Synergy 4 plate reader. The data were analyzed using Microsoft Excel and GraphPad Prism 8.

### ATPase Assay Optimization

The bioluminescence assay described above was used to determine the optimal assay conditions for the helicases. For the pH test, a 20 mM Bis-tris propane buffer was used at pH 7.0 – 9.0. For the titrations of NaCl, DMSO, MgCl_2_, ZnCl_2,_ Triton X-100, Tween 20, DTT, and TCEP, the buffer (20 mM Tris-HCl at pH 7.5) was employed. The relative activity was then evaluated by comparing these reactions to those without additives.

### Determining Kinetic Parameters with different nucleotides for 3 helicases

The optimized buffer (20 mM Tris-HCl pH 7.5, 1 mM MgCl₂, 2 mM DTT, and 0.01% Tween 20) was used in a 384-well plate format for the kinetic characterization studies of three helicases. The kinetic parameters (K_m_ and k_cat_) for each helicase/ATPase were determined using substrate titrations with ATP (up to 10 μM) or nucleotides (up to 500 nM). The reactions (15 µl) were quenched by adding an equal volume (15 µl) of Kinase-Glo buffer (Cat# V6712 or V3772), which contains components that halt enzymatic activity, allowing for accurate measurement of remaining ATP levels. Subsequently, 25 µl of the mixture was transferred to a 384-well white plate. The kinetic parameters were determined using the Michaelis−Menten equation in GraphPad Prism 8.

### Z′-Factors Determination

ATP was used at concentrations of 2 µM (for DDX1) or 4 µM (for MDA5 and LGP2) and incubated with annealed 24-mer RNA, or ds-RNA in the presence or absence of 100 nM DDX1 or LGP2, or 20 nM MDA5, respectively, for 30-60 minutes at 23°C. Luminescence signals were measured as described above, and Z’ values were calculated as previously reported ^26^ .

## Results and Discussion

We previously reported a comprehensive kinetic characterization of SARS-CoV-2 nsp13 helicase activity and developed a high-throughput screening assay to identify inhibitors for antiviral drug discovery against SARS-CoV-2 ^25^. Given the lack of similar in-depth studies on human helicases, and the urgent need for discovering human helicase inhibitors, we set out to characterize three human helicases in parallel.

While the ATPase and RNA unwinding activities of human DDX1^27–28^, MDA5^29^, and LGP2 have been previously reported ^10, 30^, their kinetics properties remain poorly characterized. To address this, we developed and optimized a luminescent -based assay to investigate their substrate specificity and determine their kinetic parameters.

Helicases exhibit NTPase activities upon binding to their respective nucleotide substrates^25–27^. We tested the helicases’ NTPase activities using various nucleotide substrates, including single-stranded RNA (ssRNA), single-stranded DNA (ssDNA), and double-stranded RNA (dsRNA). A diverse panel of nucleotides was evaluated, including a 30-base polyT (ssDNA), annealed 24- mer RNA that generated blunt-ended double-stranded RNA (ds-RNA), and a 22-bp double-stranded RNA (ds-RNA) with a 25-nt 3′ overhang. Controls without DNA or RNA were also included. Our findings revealed that all three helicases exhibited full ATPase activity in the presence of annealed 24-mer RNA or dsRNA, whereas no significant ATPase activity was observed with ssDNA or in the absence of RNA (Fig. 1).

**Figure 1.**
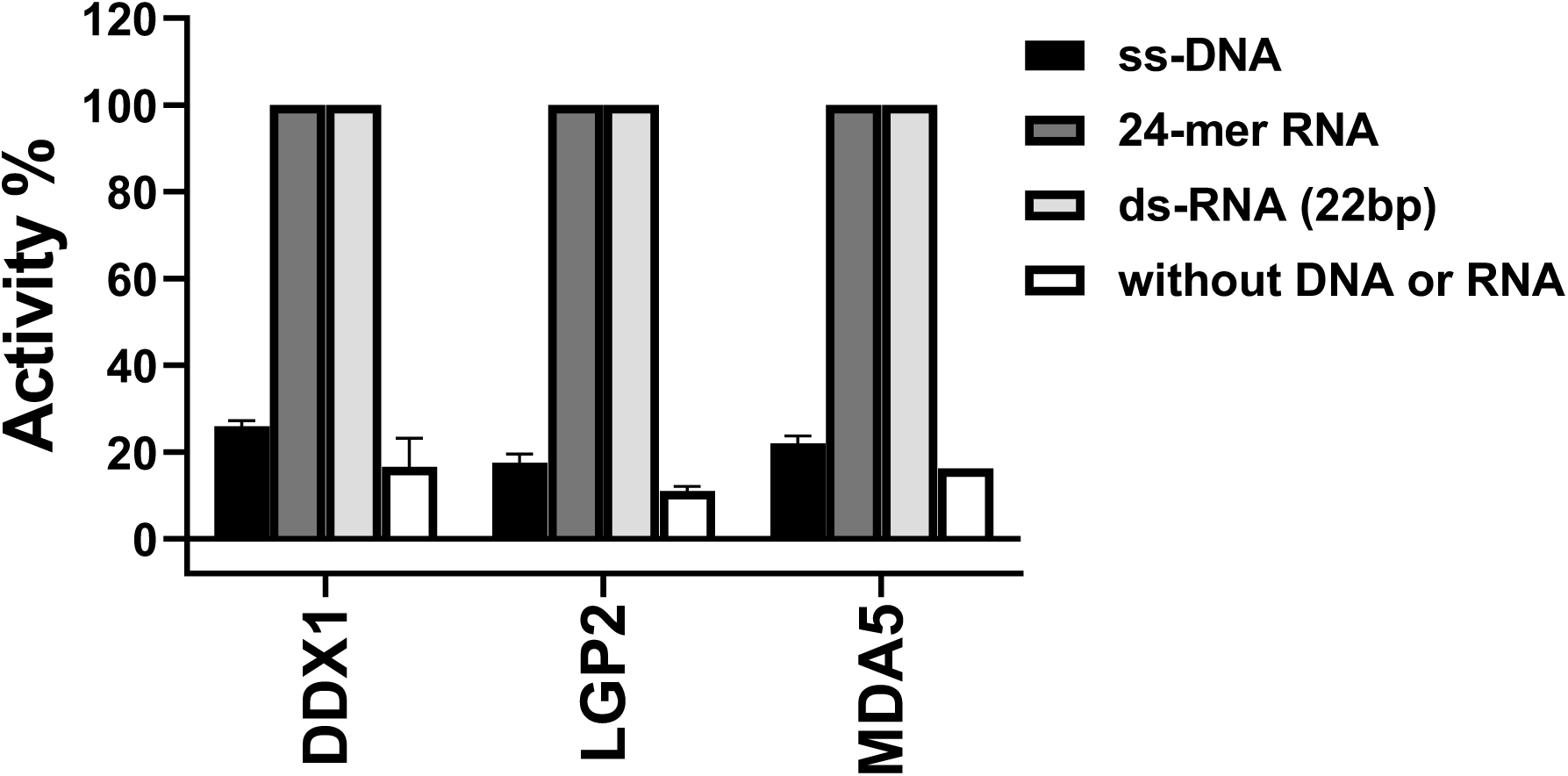
Three helicase ATPase activities were measured in the presence or absence of RNA/DNA under the conditions outlined in the ’Methods and Materials’ section. The data are presented as mean ± S.D. from three independent experiments.

The central helicase domain of MDA5 consists of Hel1, Hel2i, Hel2, and the pincer (P)^31^ responsible for RNA binding, ATP binding and hydrolysis. Additionally, the N-terminal caspase recruitment domains (CARDs, D1 and D2, 7-190 aa) mediate immune signaling^32^, while the C-terminal domain (CTD), is essential for RNA recognition and helicase activity^33^. The ATPase activities of three MDA5 constructs (306-1025 aa, 1-1025 aa and 306-890 aa) were tested in the presence of annealed 24-mer RNA and ATP. Our results demonstrated that MDA5 (306-1025 aa), which lacks the CARD domains, exhibited higher ATPase activity than the full-length MDA5 (MDA5-FL, 1-1025aa, Fig. S1). This is likely because the MDA5 (306–1025 aa) construct is more stable and remains monodisperse in solution, unlike the full-length MDA5. In contrast, the MDA5 (306-890 aa) construct, which lacks both CARDs and CTD, showed no ATPase activity (MDA5-S, 306-890aa, Fig. S1), consistent with previous findings^33^. Based on these results, all subsequent kinetic studies of MDA5 were conducted using the MDA5 (306- 1025 aa) construct.

### Assay Optimization

To optimize conditions for subsequent enzyme kinetic studies, we assessed the ATPase activity of all three helicases by testing a series of buffers and additives to determine the conditions that generate the highest signal-to-noise ratio. The various buffers tested included Tris-HCl, potassium phosphate, and HEPES, all supplemented with 1 mM MgCl₂ and 0.01% Tween 20 at pH 7.5. No significant differences were observed among them (data not shown), and Tris-HCl was selected for further experiments.

We systematically assessed the impact of pH, salts (NaCl, MgCl_2_, and ZnCl_2_), reducing agents (DTT), and other additives (DMSO, Triton X-100, Tween 20, and EDTA) on the ATPase activities of the three helicases. The ATPase activity of MDA5 remained largely unaffected across a pH range of 7.0 to 9.0 in 20 mM Bis-Tris propane buffer (Fig. 2a). Therefore, all subsequent experiments were conducted in Tris buffer at pH 7.5.

**Figure 2.**
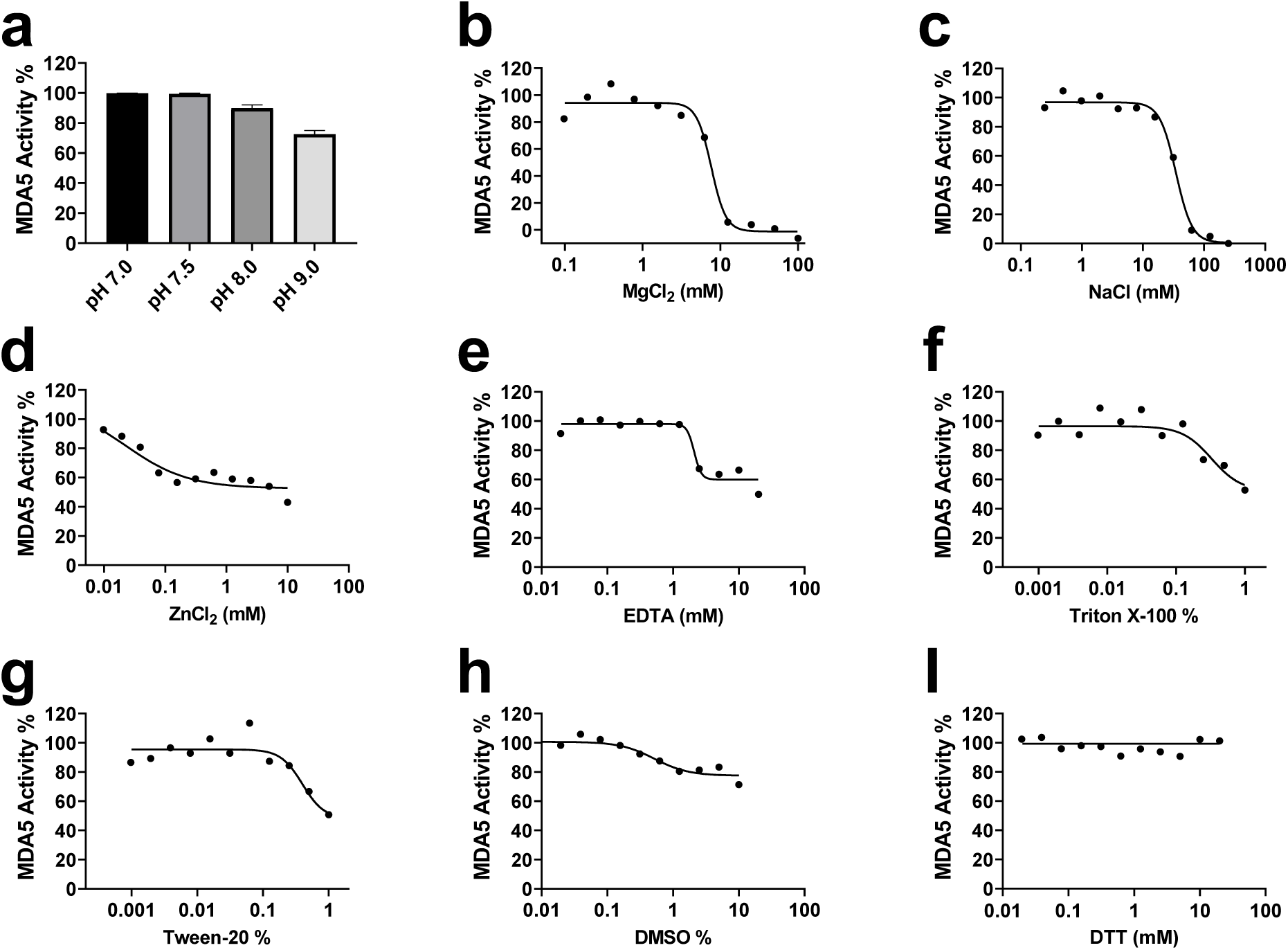
Effect of ionic strength, buffer additives pH on MDA5 ATPase activity. The ATPase activity of MDA5 in the presence of 24-mer RNA was tested as a function of additives and pH values as described under “Materials and Methods.” Data points are presented as mean ± S.D. from three experiments.

Magnesium ion (Mg^2+^) is an essential ATPase cofactor^34^; however, higher MgCl_2_ concentrations (> 10 mM) led to a significant reduction in enzymatic activity. Specifically, MDA5 activity was completely abolished (Fig. 2b), while DDX1 (Fig. S2a) and LGP2 (Fig. S3a) activities decreased by more than 50%. This inhibition may result from disruption of the optimal enzymatic environment, impairing catalytic efficiency.

The ionic strength of the buffer also strongly influenced ATPase activity, showing a sharp decline with increased concentrations of NaCl, ZnCl_2_, and EDTA (Fig. 2c-e, Fig. S2b-d, Fig. S3b-c). Specifically, DDX1 activity was completely abolished at NaCl concentrations above 25 mM (Fig. S2b), while MDA5 and LGP2 activities were lost at 62 mM (Fig. 2c) and 125 mM (Fig. S3b), respectively. Additionally, ZnCl_2_ concentrations exceeding 0.1 mM or EDTA concentrations above 2.5 mM significantly impaired the ATPase activity of all three helicases (Fig. 2 d-e, Fig. S2c-d, Fig. S3c-d). This inhibition is likely due to electrostatic interference between the enzyme and its substrate, disrupting enzymatic function.

At low concentrations (0.002-0.06%), detergents Triton X-100 or Tween 20 improved the signal-to-noise ratio. However, at higher concentrations (> 0.1%), ATPase activities declined (Fig. 2f-g, Fig. S2e-f, and Fig. S3e-f), similar to the effect observed with excessive Mg^2+,^

DMSO, commonly used as a solvent in compound libraries, had minimal impact on ATPase activity at concentrations up to 5% (Fig. 2h, Fig. S2g, and Fig. S3g). Likewise, DTT had negligible effects at concentrations up to 20 mM (Fig. 2i, Fig. S2h, and Fig. S3h). Based on these findings, all kinetic experiments were conducted under the following optimized conditions: 20 mM Tris-HCl, pH 7.5, 0.01% Tween 20, 1 mM MgCl_2_ and 2 mM DTT.

### Determination of Kinetic Parameters in the Presence of Nucleotides

The ATPase activities of three human helicases (MDA5, LGP2, and DDX1) were determined in the presence of nucleotides (annealed 24mer-RNA or ds-RNA), as described in the *Materials and Methods* section. Since ss-DNA did not significantly enhance ATPase activity of any of the three helicases, the kinetic parameters were not determined for ss-DNA. ATPase activities for MDA5A, DDX1, and LGP2 were assessed by measuring nucleotide-stimulated ATPase activity using annealed 24mer-RNA or ds-RNA and ATP as substrates.

Using optimized luminescence assay conditions, we assessed the linearity ATPase activity of all three helicases across various enzyme concentration in the presence of nucleotides (Fig. S4). Additionally, we examined reaction linearity over time at different ATP concentrations (Fig. S5). Based on these analyses, we used 20 nM MDA5 for kinetic parameter determination.

MDA5 exhibits similar ATP K_m_^app^ values with 24 mer RNA (5244 ± 484 nM) and ds-RNA (4651 ± 1008 nM). All k_cat_ values were determined based on the V_max_ observed at different ATP concentrations. The apparent k_cat_ values for MDA5 were also comparable: 210 ± 13 h⁻¹ for 24- mer RNA and 209 ± 34 h⁻¹ for ds-RNA, respectively (Table 1, Fig. 3 a - b). The K_m_ values for 24-mer RNA and ds-RNA were 2.6 ± 0.7 nM and 0.54 ± 0.02 nM, respectively (Table 1, Fig. 4a-b). Interestingly, MDA5 ATPase activity peaked at 2 – 6 nM of ds RNA, while lower or higher concentrations of ds-RNA significantly reduced activity. This may be due to several factors, including optimal binding affinity, substrate inhibition, or conformational dynamics, emphasizing the importance of both MDA5 and ds-RNA concentrations in determining overall activity in the assay. Notably, nucleotides function not only as substrates but also as activity stimulators for ATPases. Similar to the k_cat_ values, the catalytic efficiency of MDA5 for ATP (k_cat_/K ^app^) was comparable between 24-mer RNA (41.7 ± 2.7 h⁻¹ µM⁻¹) and ds-RNA (45.3 ± 3.6 h⁻¹ µM⁻¹) (Table 1, Fig. 3 a-b) in terms of ATPase activities.

**Figure 3.**
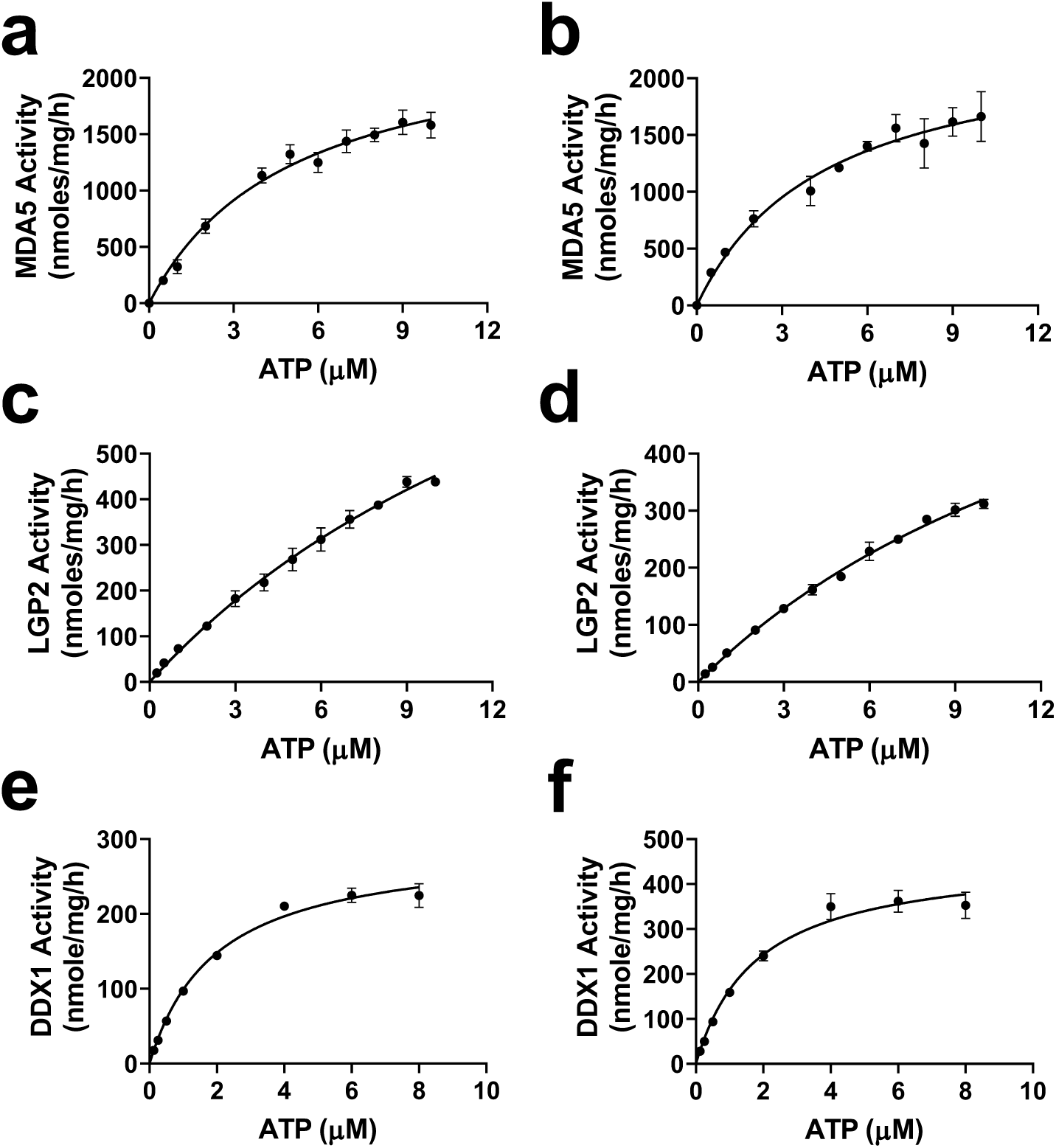
Kinetic characterization of the ATPase activities of three human helicases using varying concentrations of ATP and fixed concentrations of RNA. (a, c, and e) K_m_ determinations for ATP of the ATPase activities of MDA5 (a), LGP2 (c), and DDX1 (e) in the presence of 24-mer RNA; (b, d, and f) K_m_ determinations for ATP of MDA5 (b), LGP2 (d), and DDX1 (f) in the presence of ds-RNA. The calculated kinetic parameters are presented in Table 1. All experiments were performed in triplicate (n = 3).

**Figure 4.**
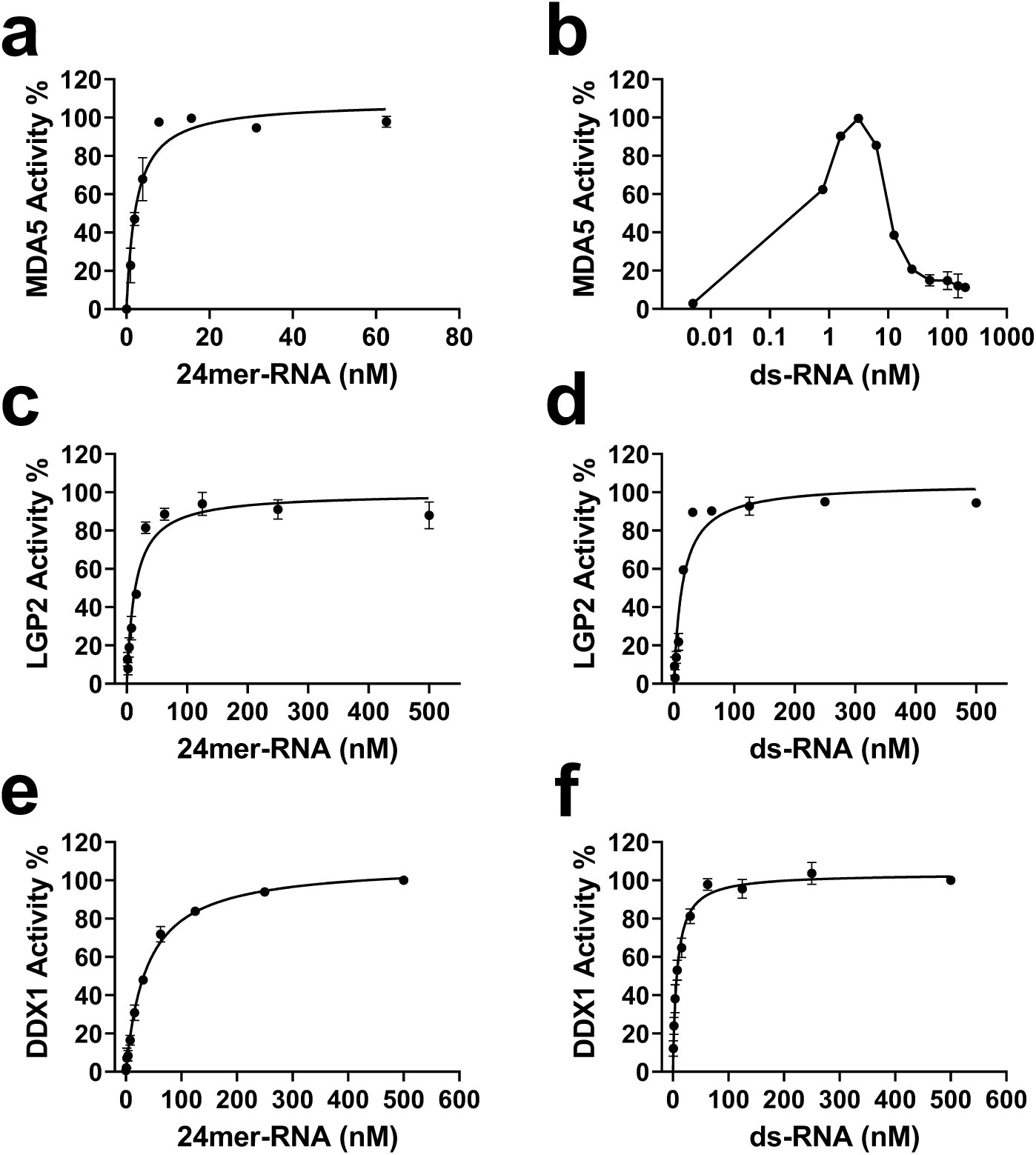
Kinetic characterization of the ATPase activities of three human helicases using varying concentrations of RNA and fixed concentrations of ATP. (a, c, e) K_m_ determinations for 24-mer RNA with MDA5 (a), LGP2 (c), and DDX1 (e), (b, d, f) K_m_ determinations for ds-RNA with MDA5 (b), LGP2 (d), and DDX1 (f). The calculated kinetic parameters are presented in Table 1. The experiments were performed in triplicate (n = 3).

**Table 1.** Kinetic Parameters for three helicase ATPase Activities.

| Helicase | Activity | Substrate | $K_m^{app}$ (nM) | $k_{cat}$ (h<sup>-1</sup>) | $k_{cat}/K_m-ATP^{app}$ (h<sup>-1</sup> μM<sup>-1</sup>) |
| --- | --- | --- | --- | --- | --- |
| **MDA5** | 24-mer RNA ATPase | ATP | 5244 ± 484 | 210 ± 13 | 41.7 ± 2.7 |
|  |  | 24-mer RNA | 2.6 ± 0.7 |  |  |
|  | ds-RNA ATPase | ATP | 4651 ± 1008 | 209 ± 34 | 45.3 ± 3.6 |
|  |  | ds-RNA | 0.54 ± 0.02 |  |  |
| **LGP2** | 24-mer RNA ATPase | ATP | > 10000 | 35.1 ± 0.5* | < 3.5 ± 0.1 |
|  |  | 24-mer RNA | 14.1 ± 1.6 |  |  |
|  | ds-RNA ATPase | ATP | > 10000 | 24.9 ± 0.6* | < 2.5 ± 0.1 |
|  |  | ds-RNA | 13.0 ± 1.0 |  |  |
| **DDX1** | 24-mer RNA ATPase | ATP | 1973 ± 207 | 19 ± 2 | 9.7 ± 0.3 |
|  |  | 24-mer RNA | 35 ± 3 |  |  |
|  | ds-RNA ATPase | ATP | 1805 ± 194 | 30 ± 3 | 16.7 ± 0.5 |
|  |  | ds-RNA | 8 ± 2 |  |  |

\*: $k_{cat}$ values were estimated based on the highest activity observed within the tested ATP concentration range, as LGP2 activity did not saturate even at 10 μM ATP.

LGP2, another member of the RIG-I-like DExH family, exhibited different catalytic behavior compared to MDA5. The ATP affinity was relatively low, with K_m_ values exceeding 10 µM (Fig. 3 c-d, Table 1). Additionally, the ATPase activity was 6 to 8-fold lower, and the catalytic efficiency was 12 to 18-fold lower than that of MDA5 (Fig. 3, Table 1). LGP2 displayed slightly higher activity with 24-mer RNA (k_cat_ value of 35.1 ± 0.5 h⁻¹) than with ds-RNA (k_cat_ value of 24.9 ± 0.6 h⁻¹, Table 1). This finding is consistent with a recent study indicating that LGP2 preferentially binds to blunt-ended ds-RNAs over RNA overhangs ^35^. However, substrate affinities were similar, with Km values of 13.0 nM for ds-RNA and 14.1 nM for 24-mer RNA (Fig. 4 c-d, Table 1).

Compared to MDA5, DDX1 has approximately 2-fold higher ATP affinity, with K_m_ ^app^ values of 1973 ± 207 nM for 24-mer RNA and 1805 ± 194 nM for ds-RNA (Table 1, Fig. 3 e-f). However, the ATPase activity (k_cat_^app^) was lower by about 7 to 10-fold, measuring 19 ± 2 h⁻¹ for 24-mer RNA and 30 ± 3 h⁻¹ for ds-RNA (Table 1), along with lower catalytic efficiency (9.7 µM⁻¹ h⁻¹ for 24mer-RNA and 16.7 µM⁻¹ h⁻¹ for ds-RNA, Table 1, Fig. 3 e-f). Additionally, ds-RNA exhibited a higher affinity (K_m_ ^app^ = 8 nM) compared to 24-mer RNA (K_m_ ^app^ = 35 nM) (Fig. 4 e-f, Table 1), and stimulated DDX1 activity approximately 1.6-fold more than 24-mer RNA (Table 1). This aligns with a previous study showing that DDX1 ATPase activity is enhanced by poly RNAs but not by DNA ^28^. However, recent studies suggest that DDX1 can hydrolyze ATP in the presence of both RNA and single-stranded DNA, though ssDNA activity remains lower than that observed with RNA^21^.

Despite sharing conserved DExD/H-box motifs essential for ATP binding, hydrolysis, and nucleic acid binding ^36^, the three helicases exhibit distinct ATPase properties (Table 1). Sequence alignment of full-length or partial helicase domains revealed limited homology. Specifically, the two RIG-I-like helicases (MDA5 and LGP2) share 44% sequence identity with each other, while they show only 19% and 17% sequence identity, respectively, with DDX1 (Table 2, Fig. S7). These sequence differences contribute to unique functional characteristics, influencing their ATPase activity and substrate preferences. Such diversity allows these helicases to fulfill specialized roles in biological processes, despite their shared DExD/H-box architecture.

**Table 2.** Matrix showing the percentage identity among the sequences of the three helicases.

| Helicase | MDA5 | LGP2 | DDX1 |
| --- | --- | --- | --- |
| MDA5 | 100 | 44 | 19 |
| LGP2 | 44 | 100 | 17 |
| DDX1 | 19 | 17 | 100 |

### Z′-factor determinations

To assess assay reproducibility and robustness for high-throughput screening, we evaluated Z′- factors in a 384-well format. MDA5 ATPase assays had Z′-factors of 0.82 for 24-mer RNA (Fig. 5a) and 0.83 for ds-RNA (Fig. 5b). The optimized assays were also amenable to screening in a 384-well format for DDX1 (0.91 of Z’-factors for both 24-mer RNA and ds-RNA, respectively, Fig. S6 a-b), and LGP2 (Z’-factors of 0.83 and 0.80 for 24-mer RNA and ds-RNA, respectively, Fig. S6 c-d).

**Figure 5.**
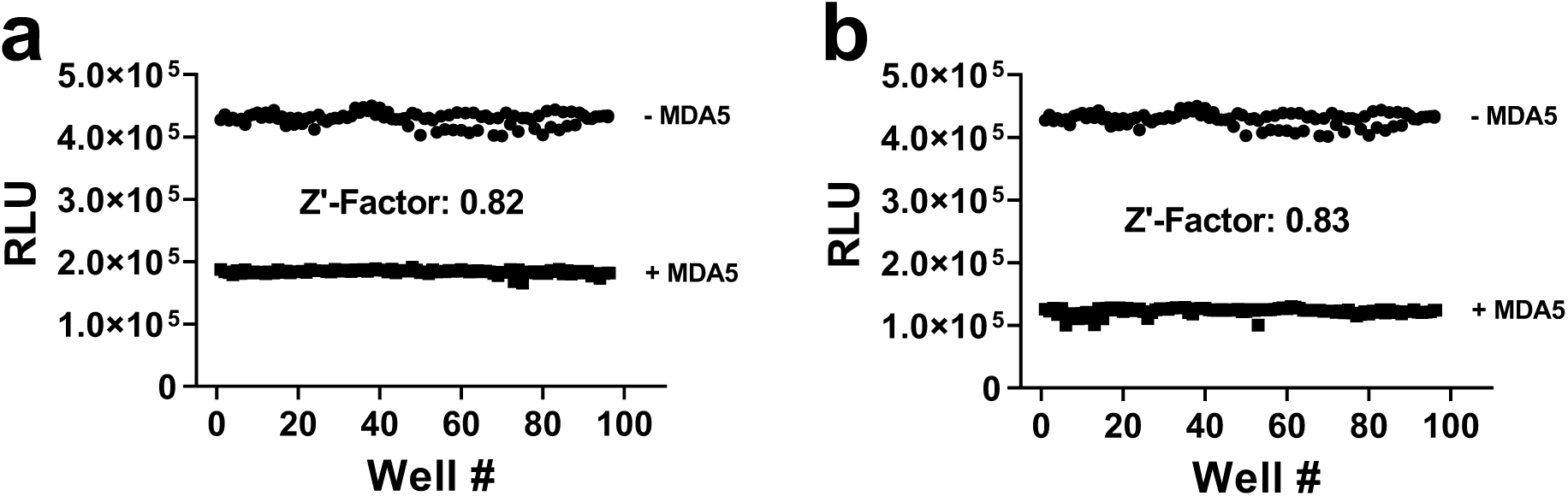
MDA5 amenability of the assays for high-throughput screening and screening in a 384- well format against a small compound library. Z’ factors for MDA5 were determined with 4 µM ATP and 30 nM 24-mer RNA (a), and with 4 µM ATP and 3 nM ds-RNA (b).

## Conclusions

The human helicases MDA5, LGP2, and DDX1 play critical roles in innate immunity and RNA metabolism. While the ATPase and helicase activities of human DExD/H-box RNA helicases have been previously reported^29^ ^21^ ^35^ ^37^, their activities and substrate specificities have not been fully characterized.

In this study, we conducted a kinetic characterization of MDA5, LGP2, and DDX1, demonstrating significant activity in the presence of annealed 24-mer RNA or dsRNA with a 3ʹ overhang. Given the robustness and cost-effectiveness of these assays, they can be readily employed as primary assays for screening large numbers of small molecules or as secondary assays for hit confirmation in other screening platforms. These assays also offer a valuable resource for evaluating computationally identified potential inhibitors of DExD/H-box RNA helicases, providing a reliable tool for advancing drug discovery and therapeutic development.

## Acknowledgements

Work reported in this report was supported in part by the Emory-Sage-SGC-JAX TREAT-AD center established by the National Institutes of Aging (NIA) under award number U54AG06518. The Structural Genomics Consortium is a registered charity (no. 1097737) that receives funds from Bayer AG, Boehringer Ingelheim, Bristol Myers Squibb, Genentech, Genome Canada through Ontario Genomics Institute [OGI-196], EU/EFPIA/OICR/McGill/KTH/Diamond Innovative Medicines Initiative 2 Joint Undertaking [EUbOPEN grant 875510], Janssen, Merck KGaA (aka EMD in Canada and US), Pfizer, and Takeda.

## Author Contributions

YL: Cloning; AS: protein expression; SDT, JGP, UHC: Protein purification; FL, IC: Biophysical assays; LH: Supervision, review and editing; FL, LH, UHC: Writing and Editing.

## Supporting Figures

**Supplementary Figure S1.**
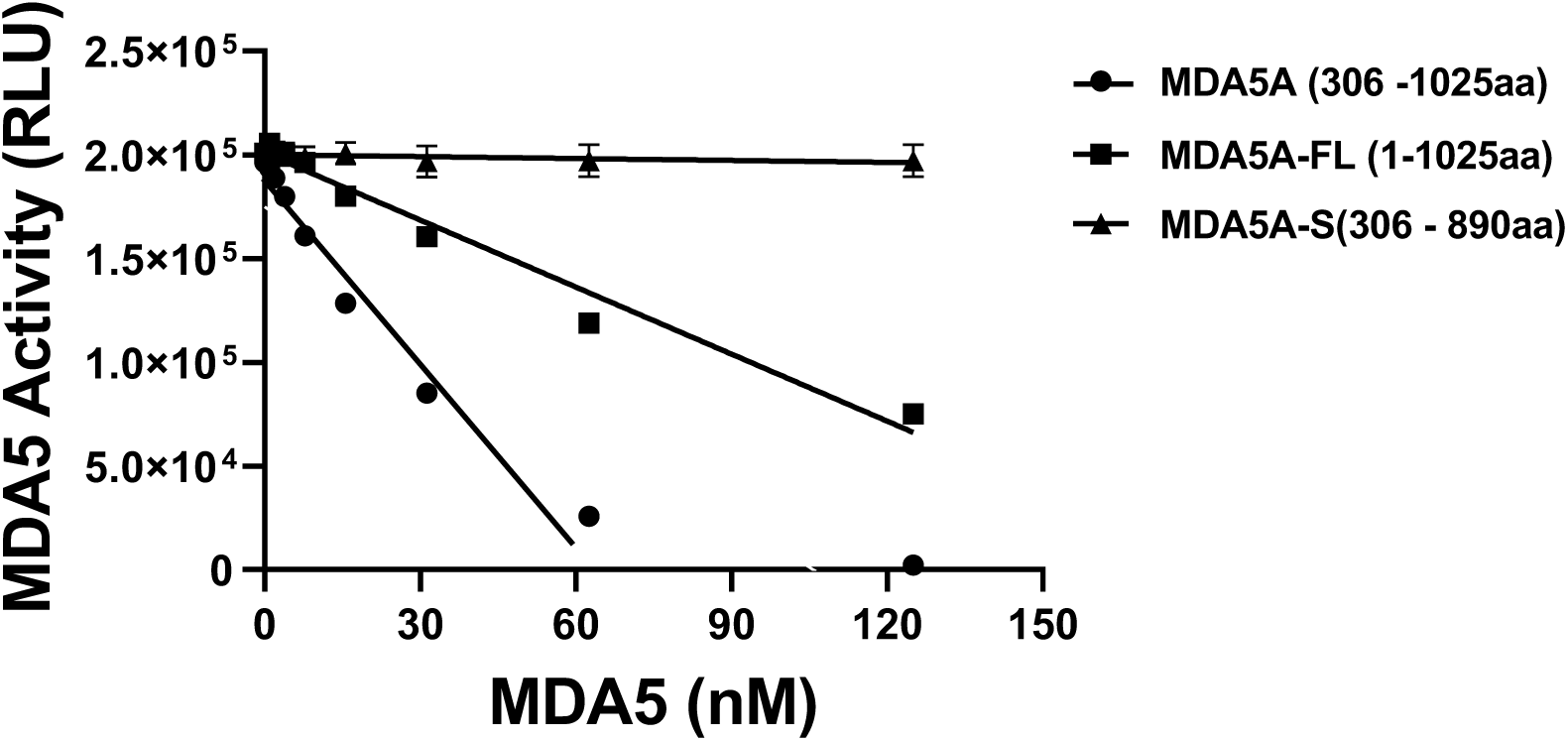
ATPase activity of different lengths of MDA5: deleted 1-306aa (●), full-length (▪) and truncated 1-306aa and 891-1025aa (▲). The experiments were conducted in triplicate (n=3).

**Supplementary Figure S2.**
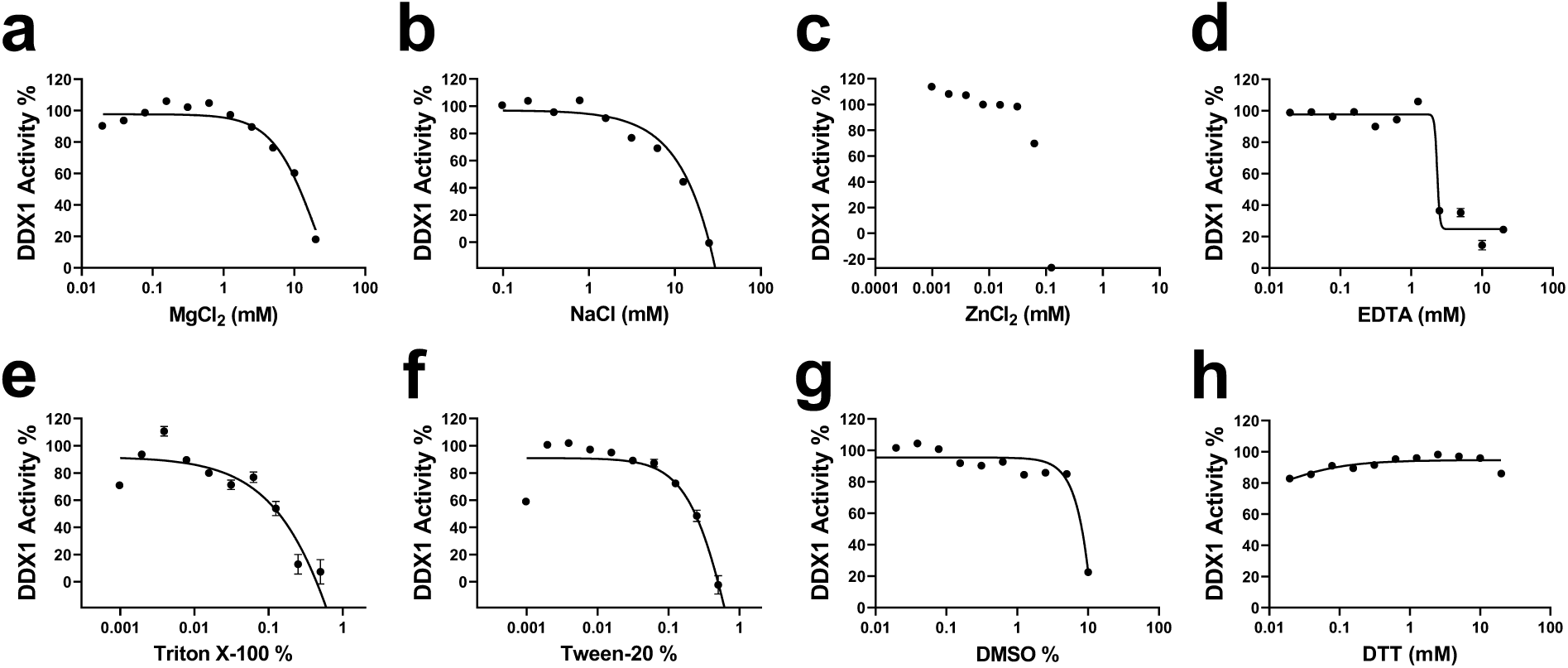
Effect of ionic strength, and buffer additives on DDX1 ATPase activity. The ATPase activity of DDX1 in the presence of 24-mer RNA was determined as a function of MgCl_2_ (a), NaCl (b), ZnCl_2_ (c), EDTA (d), Triton x-100 (e), Tween 20 (f), DMSO (g), and DTT (h) as described under “Methods and Materials”. Data points are presented as mean ±S.D. from three experiments.

**Supplementary Figure S3.**
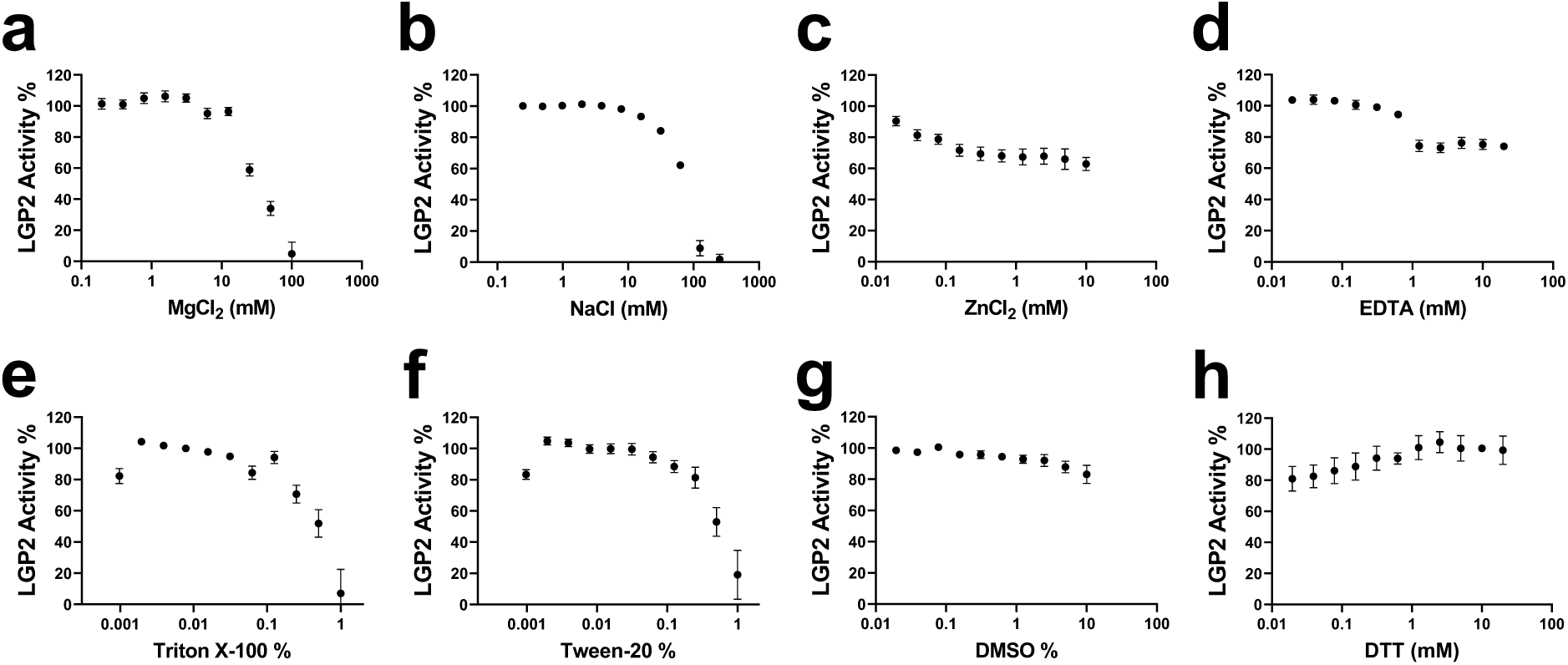
Effect of ionic strength, and buffer additives on LGP2 ATPase activity. The ATPase activity of LGP2 in the presence of 24 mer RNA was determined as a function of MgCl_2_ (a), NaCl (b), ZnCl_2_ (c), EDTA (d), Triton x-100 (e), Tween 20 (f), DMSO (g), and DTT (h) as described under “Methods and Materials”. Data points are presented as mean ±S.D. from three experiments.

**Supplementary Fig. S4.**
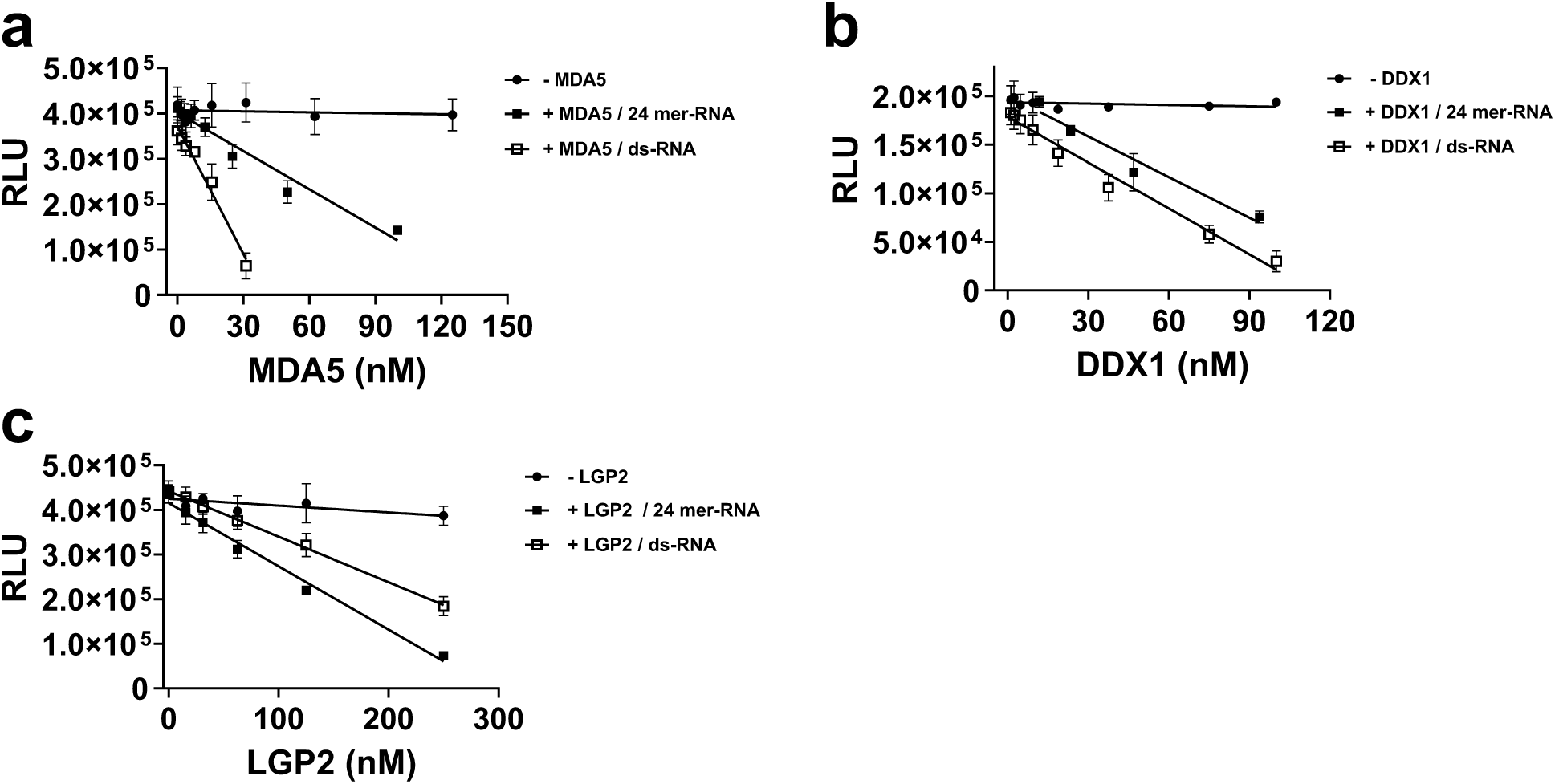
**The linearity of ATPase activities for the three helicases was tested at ATP concentrations near their K_m_ values in the presence of 24mer-RNA.** (a) MDA5 activity with varying concentrations or absence of MDA5 (●: 4.0 μM ATP and 24-mer RNA without MDA5; ▪: 4.0 μM ATP and 24-mer RNA with different concentrations of MDA5; □: 4.0 μM ATP and 24-mer RNA with different concentrations of MDA5). (b) DDX1 activity with varying concentrations or absence of DDX1 (●: 2.0 μM ATP and 24mer-RNA without DDX1; : 2.0 μM ATP and 24-mer RNA with different concentrations of DDX1; □: 2.0 μM ATP and 24-mer RNA with different concentrations of DDX1). (c) LGP2 activity with varying concentrations or absence of LGP2 (●: 4.0 μM ATP and 24-mer RNA without LGP2; ▪: 4.0 μM ATP and 24-mer RNA with different concentrations of LGP2; □: 4.0 μM ATP and 24-mer RNA with different concentrations of LGP2). The experiments were performed in triplicate (n=3). RLU: relative light unit of Luminescence.

**Supplementary Figure S5.**
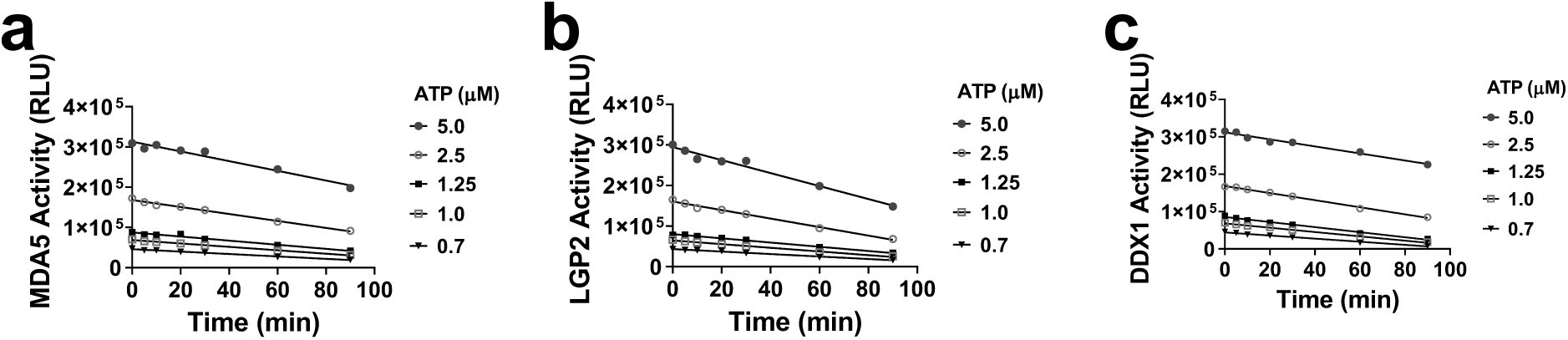
Analysis of reaction linearity over time with varying ATP concentrations. The reactions were linear up to 90 min for MDA5 (a), LGP2 (b) and DDX1 (c).

**Supplementary Figure S6.**
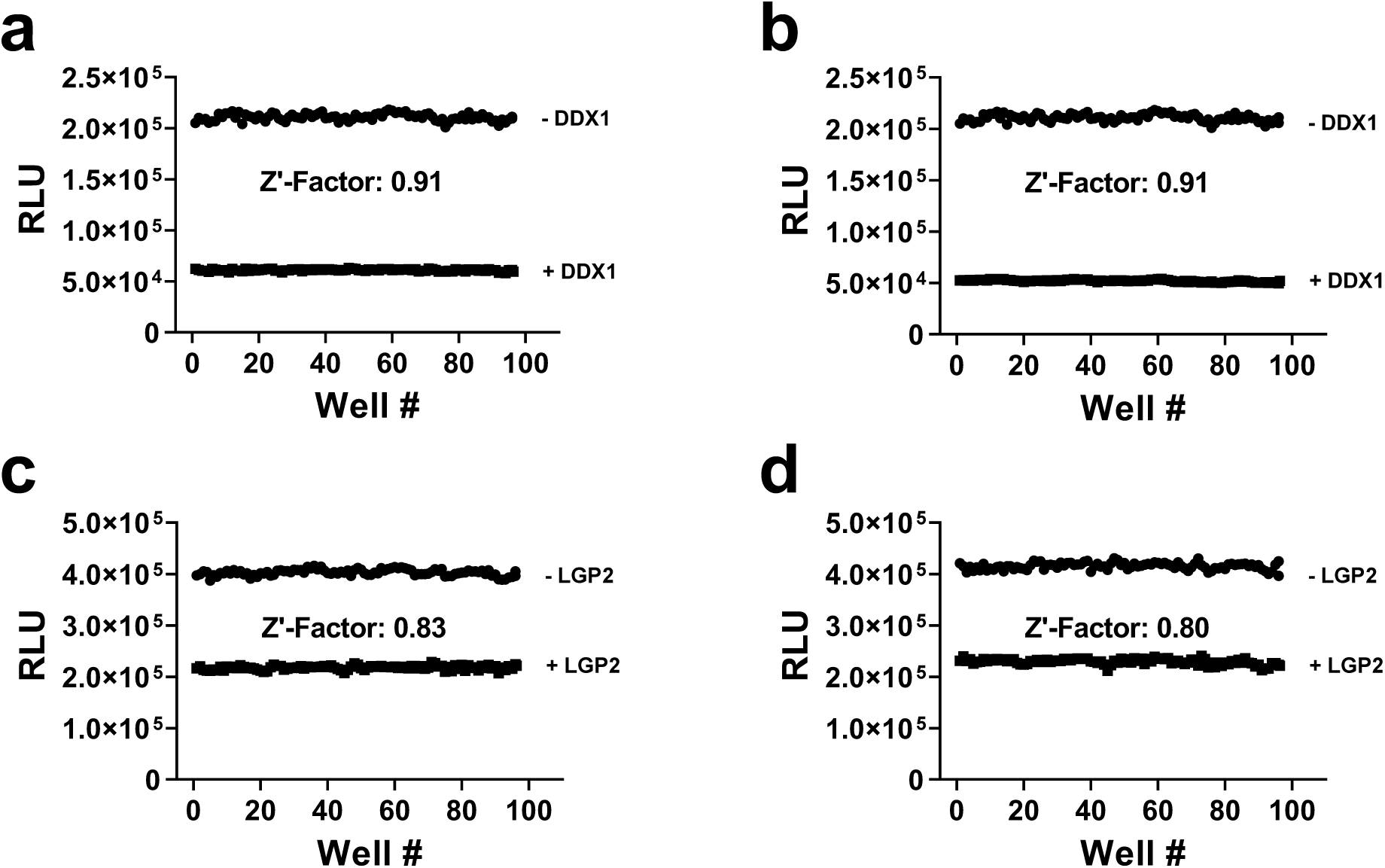
Amenability of the assays for high-throughput screening for DDX1, and LGP2. The Z’ factors were determined for DDX1 using 2 µM ATP and 100 nM 24-mer RNA (a) and 2 µM ATP with 60nM ds-RNA (b); for LGP2 using 4 µM ATP and 30 nM 24-mer RNA (c) and 4 µM ATP with 30 Nm ds-RNA (d); in the presence (▪) or absence (●) of DDX1or LGP2.

**Supplementary Figure S7.**
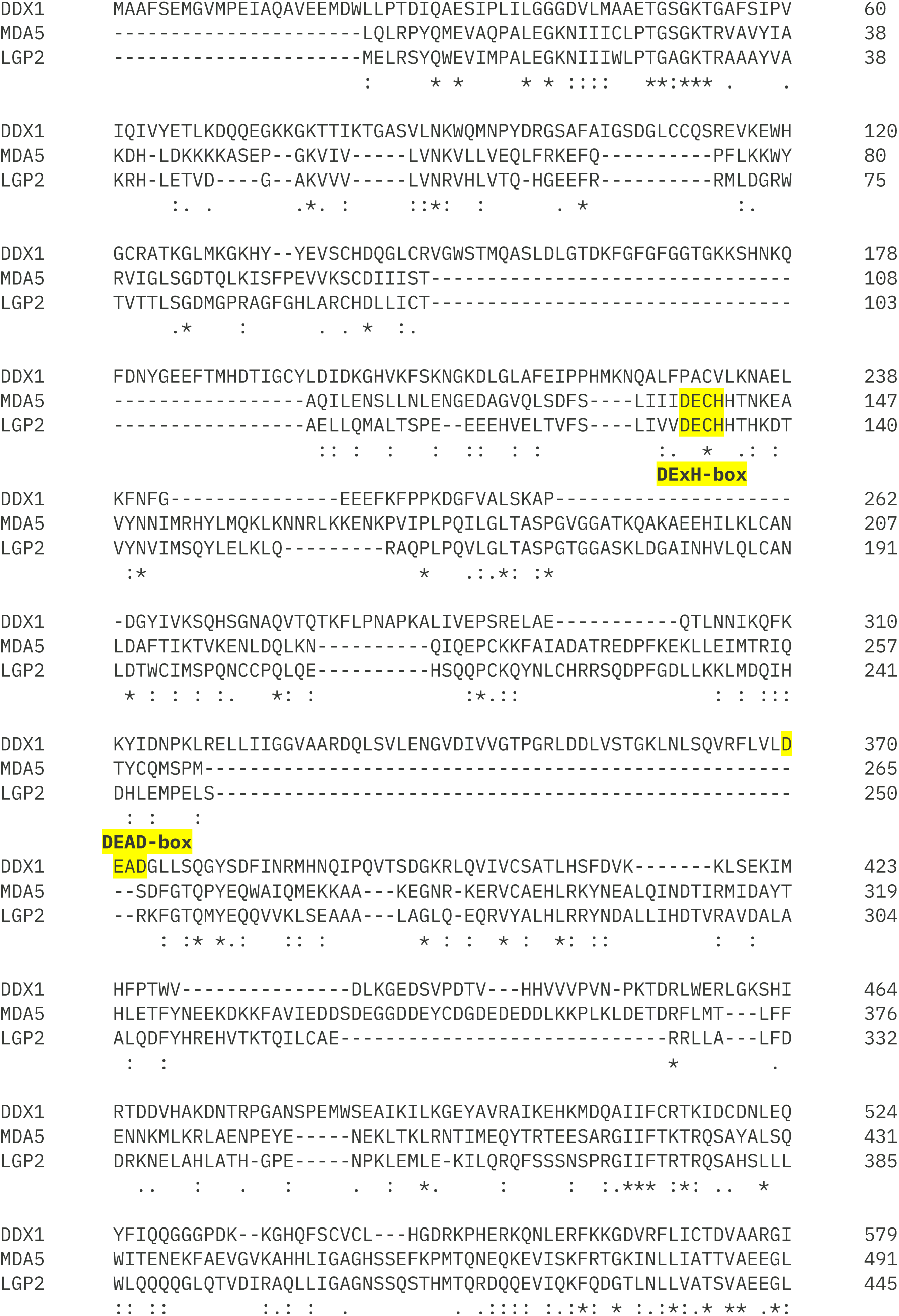
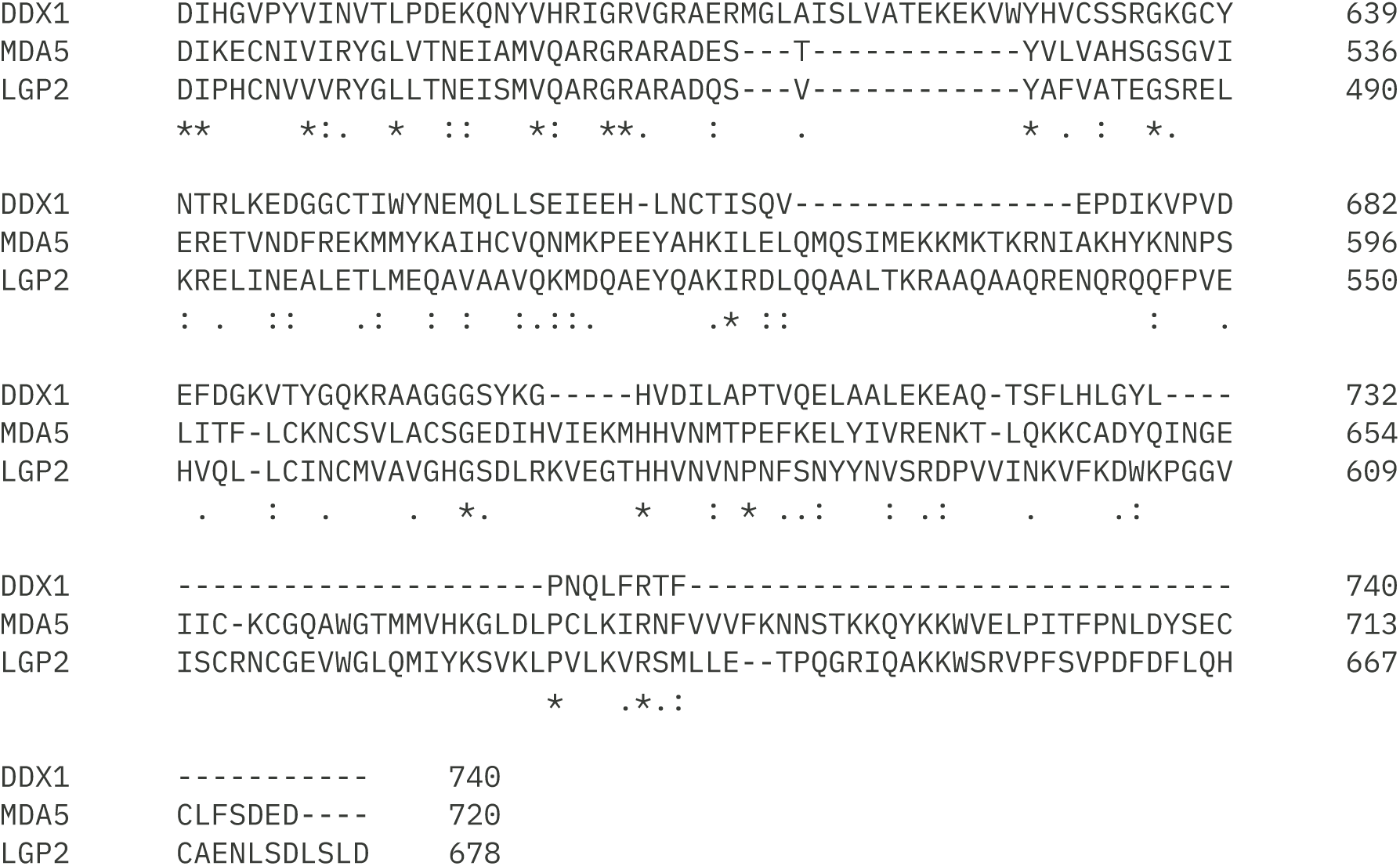
Amino acid sequence alignment of three human helicases. All amino acid sequences were obtained from the Uniprot database and were aligned using Clustal Omega program.

